# Biochemical properties and *in vitro* biological activities of extracts from seven folk medicinal plants growing wild in southern Tunisia

**DOI:** 10.1101/551515

**Authors:** Hajer Tlili, Najjaa Hanen, Abdelkerim Ben Arfa, Mohamed Neffati, Abdelbasset Boubakri, Daniela Buonocore, Maurizia Dossena, Manuela Verri, Enrico Doria

**Author notes:** These authors contributed to this study in equal measure (Hajer Tlili, Najjaa Hanen).

## Abstract

Recently, much attention has been paid to the extracts obtained from plant species in order to analyse their biological activities. Due to the climate diversity in Tunisia, the traditional pharmacopoeia consists of a wide arsenal of medicinal plant species since long used in folk medicine, in foods as spices, and in aromatherapy. Although many of these species are nearly facing extinction, only a small proportion of them have been scientifically studied. Therefore, this study explores the biochemical properties of seven spontaneous plants, which were harvested in the arid Tunisian desert: *Marrubium vulgare L., Rhus tripartita (Ucria) D.C., Thymelaea hirsute (L.) Endl., Plantago ovata Forsk., Herniaria fontanesii J. Gay., Ziziphus lotus* and *Hyoscyamus albus*. Extracts from these plants were found to contain different types of secondary metabolites (polyphenols, flavonoids, condensed tannins, crude saponins, carotenoids and alkaloids) that are involved in important biological activities. The biological activity of the extracts obtained from each Tunisian plant was assessed: first of all, leukaemia and colon cancer cell lines (K-562 and CaCo-2 respectively) were treated with different concentrations of extracts, and then the anti-proliferative activity was observed. The results showed, in particular, how the plant extract from *Rhus tripartita* significantly inhibits cell proliferation, especially on the K-562 tumour cell line. Subsequently, the anti-inflammatory activity was also assessed, and the results showed that *Herniaria fontanesii* and *Marrubium vulgare* possess the highest activity in the group of analysed plants. Finally, the greatest acetylcholinesterase inhibitory effect was exhibited by the extract obtained from *Rhus tripartita*.

In conclusion, all the Tunisian plants we analysed were shown to contain a remarkable amount of different bio-active compounds, thus confirming their involvement in several biological activities. *Rhus tripartita* and *Ziziphus lotus* were shown to be particularly effective in anti-proliferative activity, while *Herniaria fontanesii* were shown to have the best anti-inflammatory activity.

## INTRODUCTION

Nature has been a source of medicinal agents for thousands of years and an impressive number of modern drugs have been isolated from natural sources, many of them based on their use in traditional medicine. Today it is estimated that more than two thirds of the world’s population relies on plant-derived drugs; some 7,000 medicinal compounds used in the Western pharmacopoeia are derived from plants [1].

Recently, much attention has been paid to extracts and biologically active compounds isolated from plant species in order to analyse their pharmacological activities [2,3].

These plants have been used extensively in folk medicine to treat ailments and diseases [4] and are still used in the rural areas of developing countries [4,5]. In fact, the World Health Organisation (WHO) reported that around 80% of the world’s population still relies on plants as a source for primary health care [6] while traditional medicine is the only health source available for 60% of the global population [4]. Plants are the main ingredients of medicines in most traditional systems of healing and have been the source of inspiration for several major pharmaceutical drugs [7,8]. Medicinal plants are frequently the only form of cancer treatment for many people in North Africa, either due to low income or spatial distance from the urban treatment centres [9]. Tunisia has a high diversity of plants with several aromatic plant species traditionally used in folk medicines, in foods as spices, in massage and in aromatherapy. Among the 2250 species that compose Tunisia’s vascular flora [10], 1630 species are native to the arid and desert part of the country, which is characterised by low rainfall, high temperature and drying winds [11]. Remarkably, a wide range of plant species thrive under these conditions, which is of high economic and ecological significance. Due to the climate diversity in Tunisia, the traditional pharmacopoeia consists of a wide arsenal of medicinal plants. Although many of these species are nearly facing extinction [12], only a small proportion of them has been scientifically studied [8]. Therefore, this study explores and compares some biochemical and biological properties of these spontaneous plants, harvested in Tunisian arid lands.

## MATERIALS AND METHODS

### Chemicals

All reagents and standards were purchased by Sigma-Aldrich Chemicals Co. (St. Louis, MO) and Merck (Darmstadt, Germany). Cell culture media and all other supplements were purchased by ATCC® (American Type Culture Collection) Manassas, VA 20108 USA.

### Plant material

The ethno-botanical list of the plant material used in this study is reported in table 1. This list also reports the main uses of each plant in folk medicine. All the plant material was provided by the Institut des Regions Arides (IRA) in Medenine, Tunisia, where the plants *Marrubium vulgare, Herniaria fontanesii, Plantago ovata, Rhus tripartita, Thymelaea hirsuta, Ziziphus lotus* and *Hyoscyamus albus* were harvested and authenticated by botanist Dr. Mohammed Neffati according to the “Flora of Tunisia” catalogue [13]. Voucher specimens were deposited at the herbarium of the IRA.

**Table 1.** The Ethnobotanical data of studied plant species and their main uses in local communities.

| Plant species | Family | Local name | Common name | Used Part | Main uses | References |
| --- | --- | --- | --- | --- | --- | --- |
| *Herniaria fontanesii* | Caryophyllaceae | Gataba | rupturewort | Areal part | Lithiasis or diuresis<br>Treatment of kidney stones | [14]<br>[15] |
| *Hyoschymus albus* | <u>Solanaceae</u> | El bonj el abyedh | White henbane | Leaves, seeds | Sedative and antispasmodic effect<br>Pain affecting urinary tract<br>Treatment of asthma, whooping cough | [7]<br>[16] |
| *Marrubium vulgare* | <u>Lamiaceae</u> | Oum roubia | horehound | Leaves, stems | Cough and as a cholaretic in digestive and biliary com-plaints<br>Hypoglycemic activity | [17]<br>[18] |
| *Plantago ovata* | <u>Plantaginaceae</u> | Aaynem | Ispaghul | Leaves, seeds | Laxative<br>Treatment of hypercholesteremia<br>Antioxidant, anti-inflammatory | [19]<br>[20]<br>[21] |
| *Rhus tripartita* | <u>Anacardiaceae</u> | Jedari | Summaq | Roots, fruits, leaves | Diarrhea, colitis, GIT diseases, inflammatory diseases, diabetes, dysentery, hemoptysis, conjunctivitis, animal bites and poisons, hemorrhoids, sexual disease, fever, pain and various cancers | [22] [23] |
| *Thymelaea hirsuta* | Thymelaeaceae | Mithnan | Shaggy sparrow-wort | Flowers, leaves, stems | Antiseptic, anti-inflammatory, treatment of hypertension,<br>Anti-melanogenesis<br>hypoglycemic and antidiabetic | [24]<br>[25]<br>[26] |
| *Ziziphus lotus* | <u>Rhamnaceae</u> | Sidr, nbeg | Jujube | Leaves, fruits | Anti-inflammatory, analgesic and antiulcerogenic activities, liver complaints, obesity, urinary troubles, diabetes, skin infections, fever, diarrhea, insomnia | [27]<br>[28] |

### Plant extraction preparation

The aerial part of each plant was finely powdered and used for the different biochemical assays. Methanol 70% and acetone 70% extracts (0.1 g / 10 ml), macerated for 24h in shaking conditions (50 rpm), were used to assay the total content of polyphenols, the total content of flavonoids and the total antioxidant activity (DPPH test and FRAP test). For other assays (biological assays, flavonoids, condensed tannins, carotenoids, saponins and alkaloids analysis), the extraction method is described in each section.

### Total polyphenol content

The total content of phenolics was measured according to the method described by Medoua [29]. For each sample, methanol and acetone extracts (pH 2.5 using HCl) were centrifuged at 6000 rpm for 10 minutes; the supernatants were collected and the residue pellets were further washed with 1.5 ml of acetone 70% employing mechanical agitation (800 rpm, 30 minutes at 4° C) and then centrifuged. The resulting supernatants were assayed using the Folin-Ciocolteau reagent. Absorbance was measured at 725 nm and results were expressed in Gallic Acid Equivalents using a gallic acid standard curve.

### Total flavonoid content

Total flavonoid content of plant extracts was spectrophotometrically determined by the aluminium chloride method. Briefly, 150 μl of alcoholic extract, prepared as above, was mixed with 600 μl H_2_O and 45 μl 5% NaNO_2_. The solution was incubated for 5 minutes at room temperature and then 45 μl 10% AlCl_3_ was added and incubated for one more minute. Finally, 300 μl 1M NaOH and 300 μl H_2_O were added. Absorbance at 510 nm was measured from methanol and acetone extracts. Total flavonoid concentration was determined by a catechin standard curve. Results were expressed as Catechin Equivalents (CE mg 100 g-1 FW) [30]. The samples obtained using this procedure were also assayed by HPLC for quercetin and kaempferol determination.

### Condensed tannin content

Firstly, 0.5 g of each plant powder was mixed with 10 ml of acetone / methanol (containing 1% HCl) solution (7:3) and shaken (800 rpm) at 60°C for 1 hour in the dark. The samples (in triplicate) were then sonicated and centrifuged at 6000 rpm for 10 minutes and the supernatant was filtered in new test tubes. An aliquot (0.5 ml) of each extract was mixed with 3 ml of butanol:HCl (95:5, v/v) solution in screw-capped test tubes and incubated for 60 minutes at 95°C. A red coloration developed and the absorbance was then read at 550 nm. All results were expressed as mg of standard delphinidin equivalents/g dry material. A linear response was obtained between 1 μg and 5 μg of delphinidin / ml solution.

### Carotenoid content

Sample preparation was performed according to the method used by Kurilich [31] with modifications. Firstly, 0.1 g of dry plant material was added to 25 ml of a chloroform : ethanol : diethyl ether solution (2:1:0.5) containing BHT. Potassium hydroxide (1 ml, 80% w/v) was added to the mixture for saponification and the samples were stirred for 1 hour. The solution was then transferred to a separator funnel where 30 ml of a chloroform : ethanol (2:1) solution was added. After layer separation, combined organic layers were washed with 50 ml of 5% NaCl and completely dried by rotavapor. The residue was then resuspended with hexane and spectrophotometrically assayed (Perkin Elmer UV–VIS spectrophotometer) at 450 nm, using β-carotene as standard. The same samples were then used for HPLC analysis of lutein.

### Saponin content

Total saponin content (percent yield) was determined by gravimetric method as described by Kaur [32]. The methanolic extracts from each plant (1 g in 10 ml) were macerated for 24 hours and then partitioned in a water and n-butanol (1:1 ratio) solution. This solution was poured into the separator funnel and kept for 2 hours. The upper n-butanol layer was separated and the solvent was evaporated to obtain crude saponin extract.

### Alkaloid content

Total content of alkaloids was determined according to the method described by Biradar [33] with modifications. Firstly, 5 g of each sample was added to 50 ml of a solution containing 10% acetic acid in ethanol and mildly stirred for 48 hours. After filtration, the extracts were concentrated to one-quarter of the original volume and 2 ml of 3% H_2_SO_4_ and around 8 ml of water were added to reach pH 2.5. This solution was transferred to a separator funnel where 10 ml of petroleum ether: diethyl ether (1:1) solution was added. The bottom phase was collected and added to concentrate ammonium hydroxide solution until precipitation was complete (pH 8.0). The whole solution was allowed to settle and the precipitated phase was collected and washed again with ammonium hydroxide and chloroform. This phase, dried first with Na_2_SO_4_, was then completely dried by rotavapor and weighed to estimate the percentage of alkaloids.

### DPPH test

By means of the widely used 2,2-Diphenyl-1-Picrylhydrazyl (DPPH) test, it is possible to measure the anti-radical power of the prepared extracts. Different volumes of the samples (from 25 to 75 μl) were added to 1 ml of 0.2 mM DPPH solution and to a pure methanol solution for a total volume of 1.5 ml. After incubation of the samples in the dark for 60 minutes, the absorbance at 517 nm was read against a methanol control and the results were presented as EC_50_ (effective concentration, mg/ml) obtained by plotting the concentration of the tested sample with the percentage of radical scavenging activity [34].

### FRAP test

The reducing power of the extracts was determined according to the method reported by Benzie [35]. Firstly, 2.5 ml of each methanolic and/or acetone extract was added to a reaction solution with 2.5 ml of phosphate buffer (0.2 M, pH 6.6) and 2.5 ml of 1% potassium ferricyanide K_3_Fe(CN)_6_ (freshly prepared). After incubation at 50°C for 20 minutes, the mixture was centrifuged at 6000 rpm for 10 minutes and then 2.5 ml of trichloroacetic acid (10%) was added. An aliquot of 2.5 ml of the supernatant was mixed with 2.5 ml distilled water and 0.5 ml of FeCl_3_ (0.1%). The absorbance was measured at 700 nm. The EC_50_ value (mg/ml) was calculated as the effective concentration at which the reducing capacity is 50% less. Ascorbic acid was used as a reference standard.

### Anti-proliferative activity

The anti-proliferative activity of the 70% ethanol extracts (EE), macerated for 24 hours, was assessed by evaluating the cell viability by MTT assay using 3-(4,5-dim-ethylthiazol-2-yl)-2,5-diphenyl-tetrazolium bromide reagent or MTT (Sigma-Aldrich^®^) [36]. In brief, CaCo-2 cells and K-562 cells were maintained as a culture in Dulbecco’s modified Eagle’s medium in 96-well plates (2×10^5^ cells / well) and incubated at 37°C with 5% of CO_2_ for 24 hours. The medium was then replaced with another medium containing the extracts from each plant in the final concentration of 100 μg/ml. After incubation for 48 hours, this medium was replaced once again with MTT (5 mg/ml PBS)-containing medium (0.45 mg/ml final concentration). The plates were then incubated at 37°C for 48 hours. Sodium dodecyl sulphate (SDS; 10% v/v) was then added to each well (100 μl), followed by overnight incubation at 37°C. This reagent was used to solubilise and detect the formazan-crystals and its low concentration was determined by optical density. Absorbance was obtained at 570 nm using a microplate reader (Powerscan HT; Dainippon Pharmaceuticals USA Corporation, NJ, USA). Data are presented as percentage of cell viability against a control (100% of cell viability) using 100 μg/ml as concentration of the plant extracts. This concentration was chosen according to the inhibiting concentration (IC_50_) of each plant extract previously determined (data not shown).

### In vitro anti-inflammatory activity

This method was based on inhibition of albumin denaturation [37]. The reaction mixture consists of the methanolic extract, which is more effective in the extraction of the whole complex of metabolites, of each tested plant, at a concentration of 100 μg/ml (this concentration was chosen according to the inhibiting concentration (IC_50_) of each plant extract previously determined (data not shown) and 1% aqueous solution of bovine albumin fraction. The pH of the reaction mixture was adjusted using a small amount of 1N HCl. The samples were incubated at 37°C for 20 minutes and then heated at 67°C for 20 minutes. After cooling the samples, the turbidity was measured spectrophotometrically at 660 nm. The experiment was performed in triplicate. Acetyl salicylic acid (ASA) in the final concentration of 100 μg/ml was used as a reference drug and treated similarly for determination of absorbance. Percentage inhibition of protein denaturation was calculated as follows:

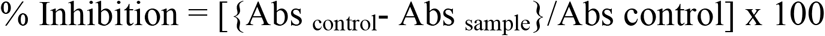

### Acetylcholinesterase inhibition

Acetylcholinesterase (AChE) enzymatic activity was measured according to the method described by Khadri [38] with some modifications. One gram of each plant material was extracted with 10 ml of 70% ethanol. After 24 hours of maceration, the samples were filtrated, dried and the residue was suspended with different volumes of water (0.0015g / ml). An aliquot of 105 μl of Tris–HCl buffer (50 mM, pH 8), 35 μl of each sample in the different concentrations and 10 μl acetylcholinesterase solution (0.26 U/ml) were mixed in 96 well plates and incubated for 15 minutes. Afterwards, 25 μl of AchI (acetylcholine iodide, 0.023 mg/ml) and 142 μl of DTNB (3 mM) were added. The absorbance was read at 405 nm when the reaction reached the equilibrium (around 10 minutes). A control reaction was carried out using water instead of extract and was considered 100% activity. Inhibition, in percentage, was calculated in the following way:

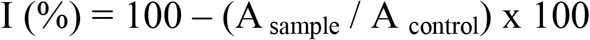

Tests were carried out in triplicate. Extract concentration providing 50% enzymatic inhibition (IC_50_) was obtained by plotting the inhibition percentage against extract concentrations.

### HPLC analysis

Chromatographic analysis of lutein was performed using a Shimadzu system equipped with DiscoVery BIO Wide Pore C18-5 column and a PDA detector (SPD-M20A). The used solvents were (A) methanol: 1M ammonium acetate 8:2 and (B) methanol:acetone 8:2. The injection volume was 20 μl and the flow rate 1ml/min. UV absorbance was settled at 450 nm. The gradient for elution was linear from 0 to 100% B in 20 minutes; after 5 minutes, 100% of A for a further 5 minutes. Finally, a linear flow of 100% A for 5 minutes was used to equilibrate the column.

### LC-ESI-MS analysis

Methanolic extracts (100 μg/ml) of plants were filtered through a 0.45 μm membrane filter before injection into the HPLC system. LC-ESI-MS analysis was performed using a LCMS-2020 quadrupole mass spectrometer (Shimadzu, Kyoto, Japan) equipped with an electrospray ionization source (ESI) and operated in negative ionization mode. Mass spectrometer was coupled online with an ultra-fast liquid chromatography system that consisted of a LC-20AD XR binary pump system, SIL-20AC XR autosampler, CTO-20AC column oven and DGU-20A 3R degasser (Shimadzu, Kyoto, Japan). A DiscoVery BIO Wide Pore C18-5 (Thermo Electron, Dreieich, Germany) (15 cm x 4.6 mm, 5 μm) was applied for analysis. The mobile phase was composed of A (0.1% formic acid in H_2_O, v/v) and B (0.1% formic acid in methanol, v/v) with a linear gradient elution: 0-14 min, 10% B; 14-24, 20% B, 27-37, 55 % B, 37-45, 100 % B, 45-50, 10% B. Re-equilibration duration was 5 minutes between individual runs. The flow rate of the mobile phase was 0.4 ml/min, the column temperature was maintained at 40°C and the injection volume was 5 μl. Spectra were monitored in mode SIM (Selected Ion Monitoring) and processed using Shimadzu LabSolutions LC-MS software.

### Statistical analysis

A descriptive analysis was performed to describe the entire results within each kind of test. Concerning the anti-proliferative activity, an unimpaired student T-test was used to compare treated cells with control cells.

Regarding the biochemical composition analysis, antioxidant activity, the in vitro anti-inflammatory activity and the acetyl cholinesterase inhibition, a one-way analysis of variance (ANOVA one-way) followed by DUNCAN test was performed to test possible significant differences among mean values from different species. The level of significance was set at P<0.05 for all analyses. Statistical analyses were performed using SPSS v.20.

## RESULTS AND DISCUSSION

### Analysis of secondary metabolites and antioxidant properties

Results about the total polyphenol content (including flavonoids and condensed tannins), the carotenoid content, the percentage of saponins and alkaloids in all the plant samples, and the antioxidant activity are presented in tables 2 and 3, respectively.

**Table 2.** Phytochemical composition of plant extracts.

|  | *Total polyphenols (mg/g)* |  | *Total flavonoids (mg/g)* |  | *Condensed tannins (mg/ml)* | *Total carotenoids (mg/ml)* | *Lutein (mg/ml)* | *Crude saponins (%)* | *Alkaloids (%)* |
| --- | --- | --- | --- | --- | --- | --- | --- | --- | --- |
|  | 70% methanol | 70% acetone | 70% methanol | 70% acetone |  |  |  |  |  |
| *H. fontanesii* | 27.23 ± 0.012<sup>d</sup> | 41.13 ± 0.050<sup>d</sup> | 8.26 ± 0.22<sup>e</sup> | 13.05 ± 0.26<sup>d</sup> | 1.31 ± 0.05<sup>f</sup> | 0.78 ± 0.09<sup>e</sup> | 0.03 ± 0.005<sup>d</sup> | 0.3 ± 0.03<sup>f</sup> | 0.3 ± 0.04<sup>d</sup> |
| *H. albus* | 22.15 ± 0.026<sup>e</sup> | 21.16 ± 0.041<sup>f</sup> | 21.57 ± 0.44<sup>b</sup> | 20.09 ± 0.40<sup>b</sup> | 2.20 ± 0.12<sup>e</sup> | 2.03 ± 0.12<sup>c</sup> | 0.19 ± 0.010<sup>b</sup> | 0.7 ± 0.08<sup>d</sup> | 0.4 ± 0.04<sup>c</sup> |
| *M. vulgare* | 18.15 ± 0.006<sup>d</sup> | 16.07 ± 0.008<sup>g</sup> | 14.46 ± 0.35<sup>d</sup> | 12.49 ± 0.29<sup>d</sup> | 4.38 ± 0.09<sup>d</sup> | 3.21 ± 0.18<sup>a</sup> | 0.43 ± 0.040<sup>a</sup> | 0.4 ± 0.03<sup>e</sup> | 0.3 ± 0.03<sup>d</sup> |
| *P. ovata* | 23.92 ± 0.021<sup>e</sup> | 27.60 ± 0.039<sup>e</sup> | 17.63 ± 0.41<sup>c</sup> | 16.39 ± 0.26<sup>c</sup> | 5.13 ± 0.11<sup>c</sup> | 2.93 ± 0.15<sup>b</sup> | 0.18 ± 0.007<sup>b</sup> | 0.8 ± 0.05<sup>c</sup> | 0.1 ± 0.01<sup>e</sup> |
| *R. tripartita* | 91.58 ± 0.049<sup>a</sup> | 103.67 ± 0.071<sup>a</sup> | 14.26 ± 0.61<sup>d</sup> | 23.51 ± 0.25<sup>b</sup> | 9.96 ± 0.22<sup>a</sup> | 0.85 ± 0.06<sup>e</sup> | 0.04 ± 0.005<sup>d</sup> | 1.2 ± 0.07<sup>a</sup> | 0.4 ± 0.03<sup>c</sup> |
| *T. hirsuta* | 47.25 ± 0.033<sup>b</sup> | 50.09 ± 0.051<sup>b</sup> | 27.60 ± 0.72<sup>a</sup> | 36.83 ± 0.31<sup>a</sup> | 7.63 ± 0.17<sup>b</sup> | 1.25 ± 0.11<sup>d</sup> | 0.10 ± 0.006<sup>c</sup> | 0.4 ± 0.05<sup>e</sup> | 1.3 ± 0.06<sup>a</sup> |
| *Z. lotus* | 31.52 ± 0.045<sup>c</sup> | 43.02 ± 0.044<sup>c</sup> | 12.32 ± 0.31<sup>c</sup> | 17.50 ± 0.25<sup>c</sup> | 2.99 ± 0.11<sup>e</sup> | 0.99 ± 0.07<sup>e</sup> | 0.03 ± 0.002<sup>d</sup> | 0.9 ± 0.10<sup>b</sup> | 0.5 ± 0.03<sup>b</sup> |

Data are presented as mean values ± standard deviation (n=3). Statistical analysis: ANOVA test and DUNCAN test. The different letters above the values in the same column indicate significant differences (p<0.05). Values with the same superscript letters in the same column are not significant.

**Table 3.** Antioxidant activity of plant extracts

|  | *DPPH* (*EC*<sub>50</sub> mg/ml) |  | *FRAP* (*EC*<sub>50</sub> mg/ml) |  |
| --- | --- | --- | --- | --- |
|  | 70% methanol | 70% acetone | 70% methanol | 70% acetone |
| *H. fontanesii* | 1.30 ± 0.023<sup>b</sup> | 1.26 ± 0.025<sup>b</sup> | 0.19 ± 0.006<sup>c</sup> | 0.12 ± 0.004<sup>d</sup> |
| *H. aureus* | 0.15 ± 0.003<sup>g</sup> | 0.22 ± 0.001<sup>e</sup> | 0.32 ± 0.003<sup>a</sup> | 0.46 ± 0.042<sup>b</sup> |
| *M. vulgare* | 1.11 ± 0.012<sup>c</sup> | 1.66 ± 0.027<sup>a</sup> | 0.08 ± 0.003<sup>g</sup> | 0.06 ± 0.008<sup>e</sup> |
| *P. ovata* | 0.56 ± 0.010<sup>d</sup> | 0.48 ± 0.004<sup>c</sup> | 0.17 ± 0.012<sup>d</sup> | 0.20 ± 0.006<sup>c</sup> |
| *R. tripartita* | 0.44 ± 0.008<sup>e</sup> | 0.30 ± 0.003<sup>d</sup> | 0.22 ± 0.010<sup>b</sup> | 0.75 ± 0.001<sup>a</sup> |
| *T. hirsuta* | 1.90 ± 0.015<sup>a</sup> | 0.17 ± 0.001<sup>f</sup> | 0.11 ± 0.005<sup>f</sup> | 0.10 ± 0.010<sup>d</sup> |
| *Z. lotus* | 0.19 ± 0.036<sup>f</sup> | 0.11 ± 0.002<sup>g</sup> | 0.13 ± 0.005<sup>e</sup> | 0.18 ± 0.006<sup>c</sup> |

Data are presented as mean values ± Standard deviation (n=3). Statistical analysis: ANOVA test and DUNCAN test (p<0.05). Values with the same superscript letters in the same column are not significant.

Antioxidant activity measured by the DPPH test and the FRAP test, in both methanol and acetone extracts, showed remarkable variability. Considering the results obtained by using both extraction methods, the highest radical scavenging activity demonstrated by the DPPH test was found in the methanol extract of *Z. lotus:* it was around 10 times higher than the average value observed in the other plant samples extracted with the same solvent. The extraction carried out with acetone provided lower EC_50_ results (hence higher antioxidant activity) than those registered for the methanol extract, except in *M. vulgare* and in *H. albus*. In particular, in *T. hirsuta*, the antiradical activity measured in the acetone extract was almost 11 times higher than that observed in the methanol extract.

The FRAP test revealed how the most significant difference in the ferric reducing antioxidant power was observed in *R. tripartita* where the methanol extract showed an antioxidant activity that was almost 3.5 times higher than the acetone extract. Regarding total polyphenol content, few significant differences were found between the two types of solvent extraction. In particular, the acetone extract of *H. fontanesii* showed a phenolic content that was around 50% higher than the methanol extract. *R. tripartita* presented the highest value of these secondary metabolites: almost 3.5 times higher than the average registered for the other plants, regardless of the solvent used for extraction. *T. hirsuta* showed the highest content of total flavonoids, both for methanol and acetone extract. There are not many data available in literature about the plants examined in this study, so making a comparison of the results we obtained is not easy. Nonetheless, a few papers reported the amount of phenolics in some of these plants; recently, in a review about the biochemical composition of different parts of *Z. lotus*, Abdoul-Azize [39] reported around 7 mg/g of phenolics to be present in the leaves, almost 5 times less than the amount measured in our study. Moreover, in the same review, the tannin content observed in the leaves was the same that we measured in the samples described in this paper (around 3.0 mg/g). Conversely, the content of phenolics in the leaves of *T. hirsuta* and *R. tripartitum*, including flavonoids, and the DPPH values found by Akrout [40] and Itidel [41] were in line with those measured in our study. When Alghazeer [42] studied the antioxidant activity of some plants growing in Libya, he found two times higher the amount of polyphenols in *H. albus* than we found in this study. The total carotenoid content in the Tunisian plants we analysed was quite variable. *M.vulgare* and *P.ovata* showed the highest content of these pigments (3.21 mg/g and 2.90 mg/g respectively), while *R. tripartita* presented the lowest value (0.85 mg/g). These data, presented in Table 2, reflect the lutein content measured in the plant leaves, of which *Marrubium* was found to contain the highest amount (0.43 mg/g): around 4.5 times higher than the average of the other plants. There are no available data in literature about the content of lutein or carotenoids in the plants examined in this work, so the results shown in the present paper represent the first indication of the level of these pigments.

Mass spectrometry analysis (table 4) revealed the presence, in the plant extracts, of a large variation of flavonoids involved in several biological activities. In particular, *R. tripartita* showed a richer profile of flavonoids than the other plants we analysed, with a high amount of luteolin-7-o-glucoside and apigenin-7-o-glucoside, which are both involved in cancer prevention [43]. *Z. lotus* showed around 15 times higher the amount of rutin, a glycoside of the flavonoid quercetin with a documented anti-inflammatory and anti-carcinogenic activity [44], than the average value found in the other Tunisian plants. In *H. fontanesii*, high concentrations were found of catechin, epicatechin and rutin, which are all molecules involved in the prevention and treatment of chronic diseases in humans such as inflammatory diseases [45]. Finally, *T. hirsuta* showed the highest amount of the flavonoid kaempferol: more than 10 times higher than the average amount registered for the other examined plants.

**Table 4.**
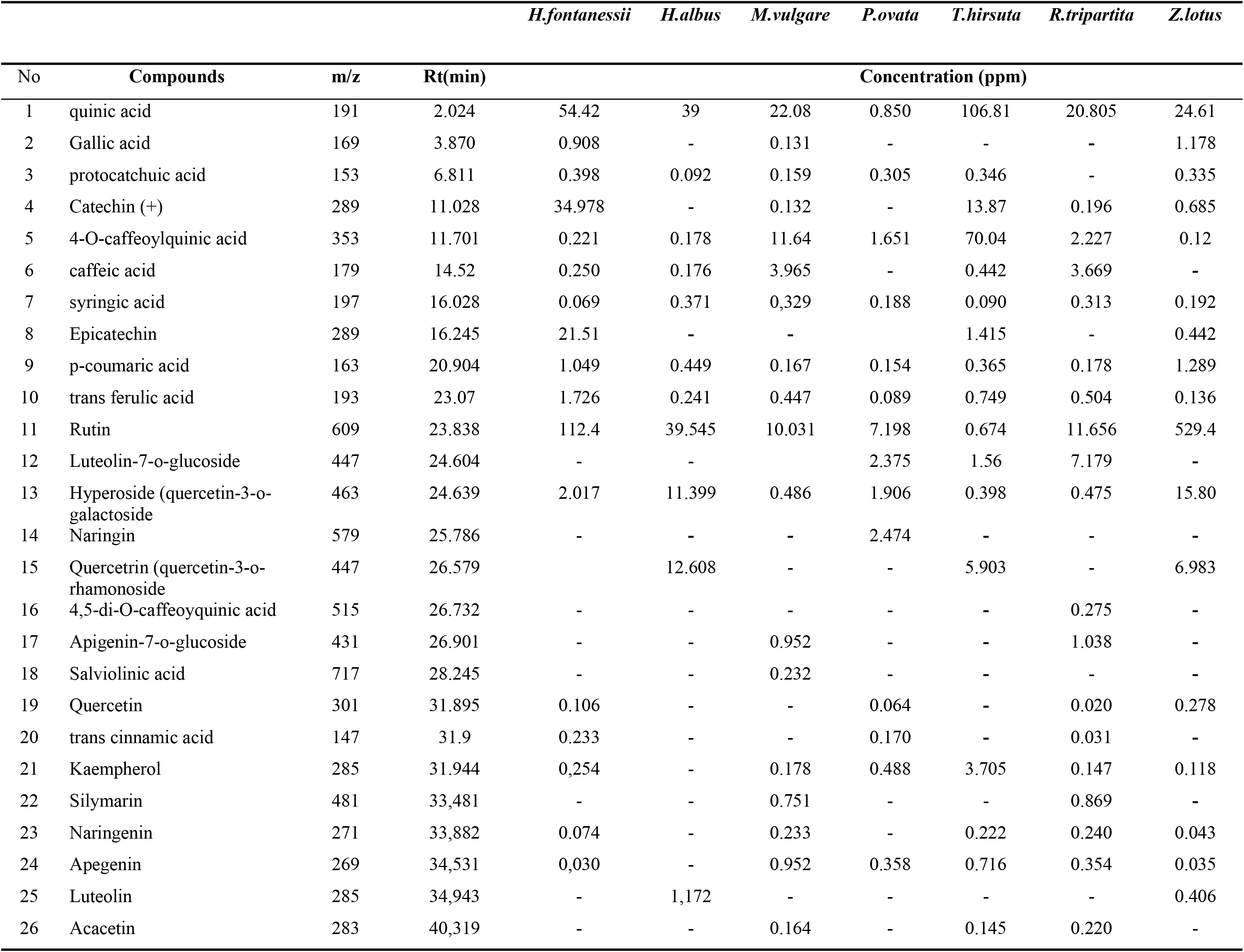
Mass spectrometry analysis of different flavonoids present in the plant alcohol extracts.

Data about the content of saponins and alkaloids in the examined aromatic plants are expressed as a % of the dry weight and are shown in Table 2. Regarding the content of the glycosides triterpenoids, *R. tripartita* showed the highest amount (12 mg/g DW), while in the other plant leaf extracts, the amount of saponins ranged between 0.3% and 0.9%, with a high sampling variance σ^2^ of 0.09. The saponin content measured in *Z. lotus* (0.9%) was more than double the amount (0.4%) found by Abdoul-Azize [39], while in *P. ovata*, the concentration of these compounds (0.8%) was in line with that observed by Mamta [46] which measured a content of 0.7% in the leaves.

The highest alkaloid percentage was registered in *T. hirsuta* (1.3%), while the lowest value was found in *P. ovata* (0.1%); in all the other plants, the values ranged between 0.3 and 0.5%. There are few data available in literature about the measurement of the total alkaloid content in these plants; the same percentage (0.3) registered for Marrubium was found by Ohtera *et al*. [47] for the betonicine and stachydrine alkaloids.

### Anti-proliferative activity

The cytotoxicity of the seven plant extracts was performed against two human cancer cell lines, CaCo-2 (colon carcinoma) and K-562 (myelogenous leukemia). Results revealed a concentration and species-dependent cytotoxic effect of the examined extracts; the cell viability of the two cancer cell lines, expressed as a percentage, is shown in Figure 1. Out of the seven plant extracts, *Rhus tripartita* was found to be the most effective in inhibiting cell proliferation (IC_50_ value < 50 μg/ml), in particular of the K-562 leukaemia tumour cell line. These data confirmed the results obtained by Najjaa *et al*. [8], who found that *Rhus tripartita* displayed the strongest anti-cancer activity against colon adenocarcinoma cell lines (DLD-1). In fact, data presented in this paper showed that *Rhus tripartita* extracts, perhaps due to the high variability and concentration of polyphenols, possess the highest growth inhibitory and cytotoxic effects on carcinoma and leukaemia cell lines.

**Fig 1.**
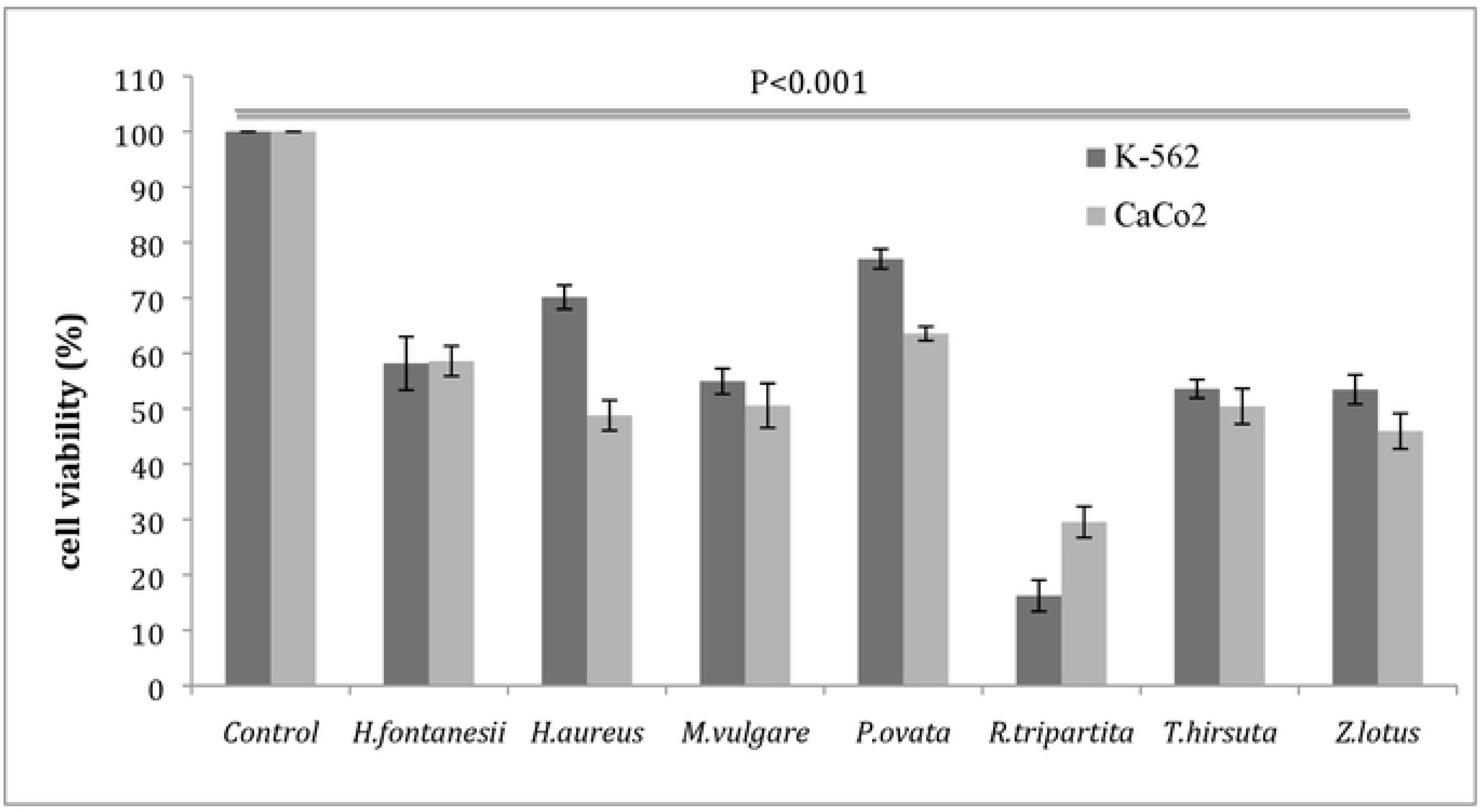
Cell viability (%). Anti-proliferative activities of the different ethanol extracts (100 μg/ml) of Tunisian medicinal plants tested on two neoplastic cell lines (K-562 and CaCo-2). Data are presented as mean values ± standard deviation (n=3). Statistical analysis: unpaired STUDENT T-test. The results were statistically significant compared with the untreated cells (control) (p <0.001).

### In-vitro anti-inflammatory activity

Denaturation of proteins, with the consequent loss of their biological activity, is a well-documented cause of inflammation [48]; therefore, agents that cause the prevention of precipitation of denatured protein aggregates and protein condensation are considered useful in disease treatment such as rheumatic disorders, cataracts and Alzheimer’s disease [49, 50]. As part of the investigation into the mechanism of anti-inflammatory activity, the ability of the examined plant extracts to inhibit protein denaturation was studied and results are presented in Figure 2.

**Fig 2.**
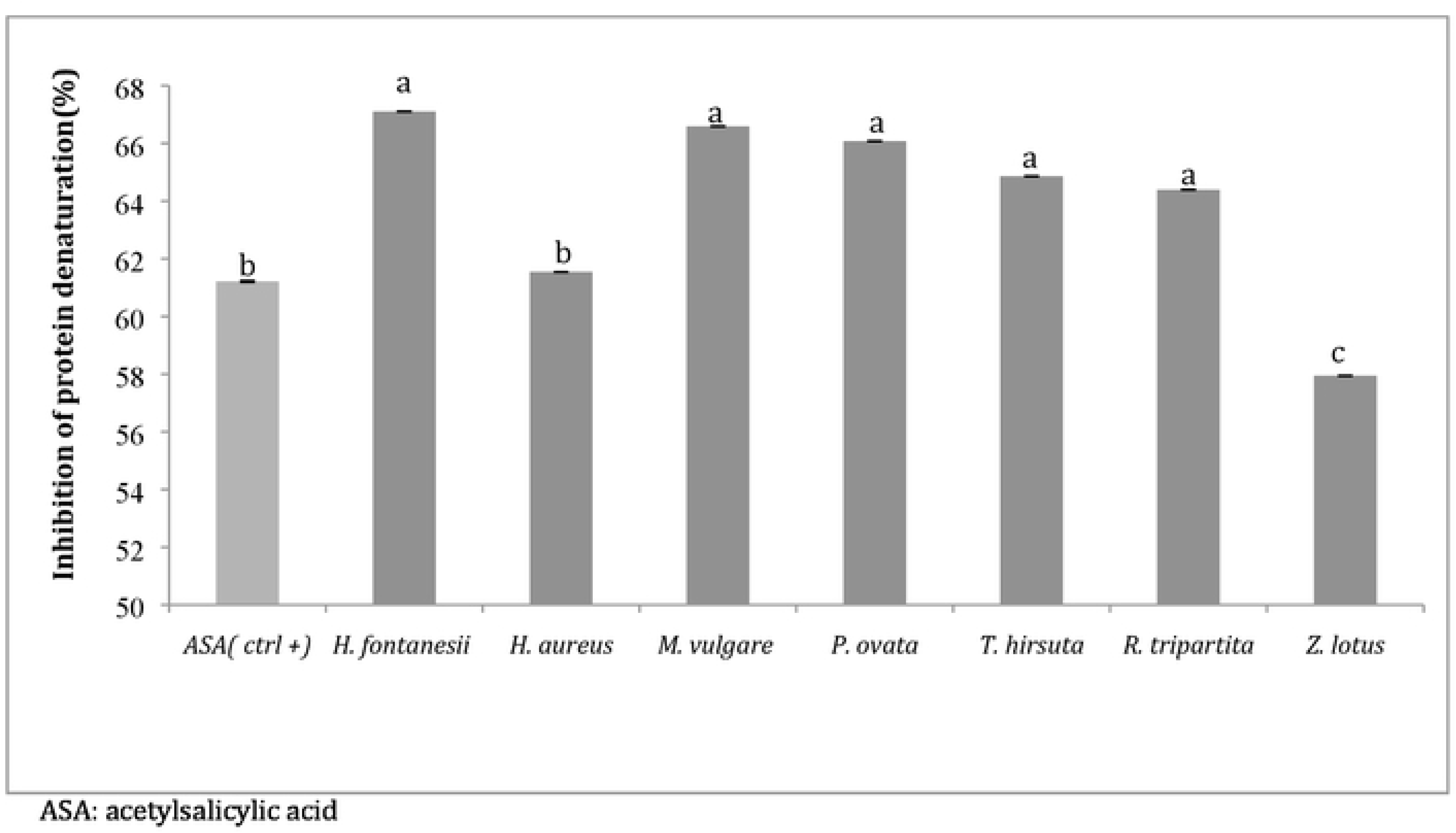
Inhibition of protein denaturation (%). Anti-inflammatory activity of the different methanolic extracts (100 μg/ml) of the tested Tunisian medicinal plants. Data are presented as mean values ± standard deviation (n=3). Statistical analysis: ANOVA test and DUNCAN test. ^a,b,c^ Different letters above the bars indicate significant differences (p<0.05).

No remarkable differences were registered among the examined plant extracts; except for *Z. lotus*, which presented a slightly lower percentage of inhibition of protein denaturation than the other aromatic plants, a concentration of 100 μg/ml of plant extract was shown to be very effective in inhibiting heat induced albumin denaturation in a range between 59.94% and 67.10%. Acetylsalicylic acid (ASA), used as a control reference, showed 61.21% inhibition.

### Acetylcholinesterase inhibition

Aromatic plant extracts were tested to determine their ability to inhibit acetylcholinesterase activity. This enzyme (AChE) regulates hydrolysis of acetylcholine (ACh) in the brain, so it is an important target for the treatment of Alzheimer’s disease (AD) [51], a feature of which is ACh deficiency.

Results, expressed as IC_50_ values and presented in Figure 3, showed that, except for *H. albus*, all the plants are able to inhibit AChE activity by 50%, at a concentration less than or equal to 1 mg/ml. This inhibitory activity could be attributed to the chemical compositions of plants mainly containing flavonoids, phenolic acids and tannins, as well as, to the possible synergistic interaction between these components [52, 53]. The results obtained in this work are in concordance with those found by Orhan et *al*. [54], who used acetone extracts from several aromatic plants, and in the range of the values reported for other Lamiaceae and Fumariaceae species [55, 56].

**Fig 3.**
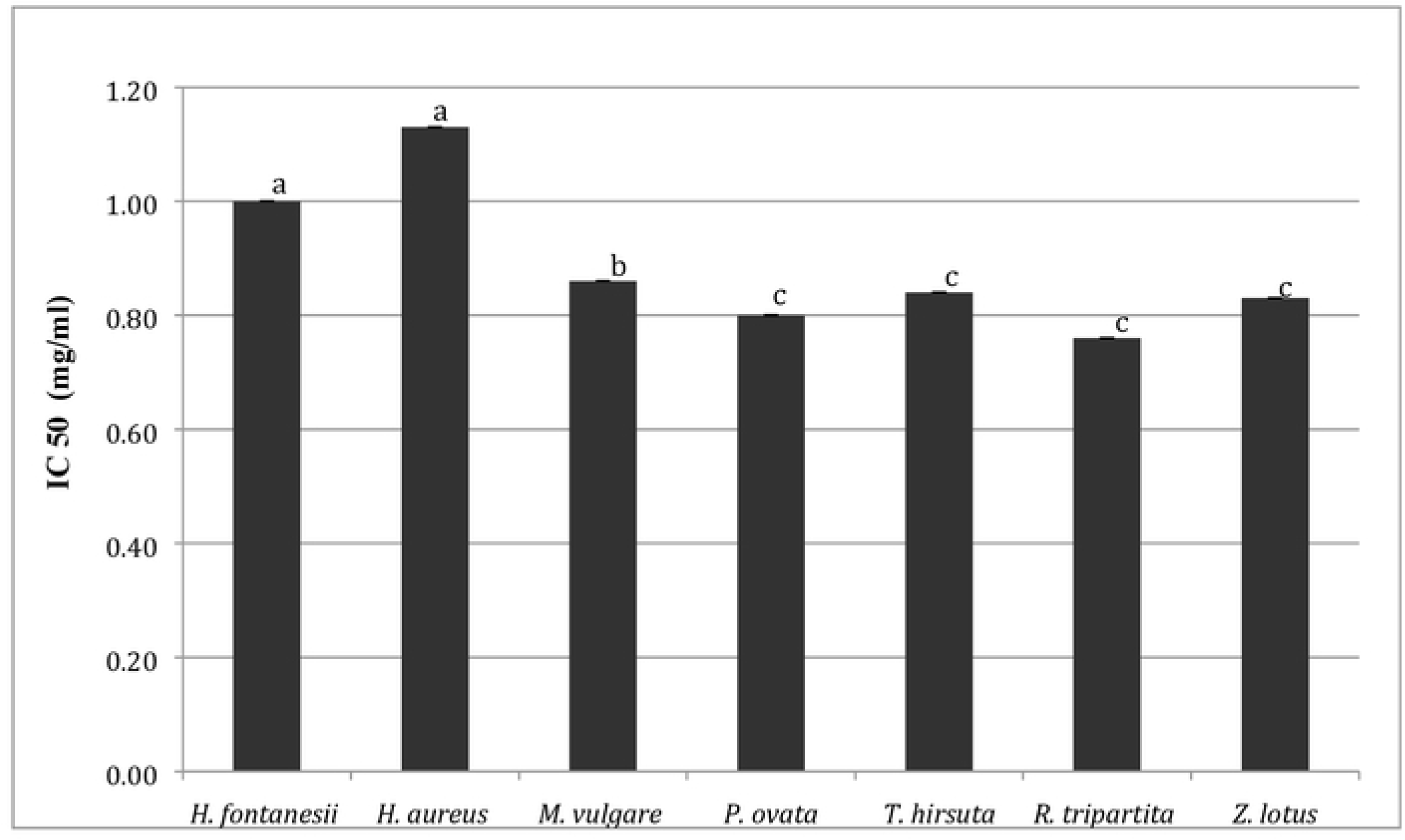
IC_50_ (mg/ml). Acetylcholinesterase activity inhibition of ethanol extracts of the tested Tunisian medicinal plants. Data are presented as mean values ± standard deviation (n=3). Statistical analysis: ANOVA test and DUNCAN test. ^a,b,c^ Different letters above the bars indicate significant differences (p<0.05).

## CONCLUSIONS

This work offers an overview of some biochemical and biological properties of seven aromatic plants, traditionally used in the Tunisian region in folk medicine. Both biochemical and biological tests were performed to provide a complete framework for each plant examined in this study. All the tested Tunisian plants showed a remarkable presence of secondary metabolites, involved in several biological activities. In particular, *Rhus tripartita* has a high content of polyphenols and saponins, responsible for the significant anti-proliferative activity. Due to the abundance of bioactive metabolites, all the extracts obtained by the plants were shown to be able to inhibit AChE activity by 50%, at a concentration less than or equal to 1 mg/ml; moreover, these extracts were shown to be efficient, with the exception of *Z. lotus*, in the prevention of precipitation of the denatured protein aggregates involved in inflammation. In conclusion, all the data confirm the importance of the Tunisian local vegetation as a potential source of various bioactive phytochemical compounds; the investigation is based on the need for different biological agents from natural sources with potent activity and lesser side effects as substitutes for chemical therapeutics.

## Acknowledgements

This work was conducted within an Agreement between the University of Pavia, Italy and the Range Ecology Laboratory, Arid Lands Institute of Medenine, Tunisia. The authors thank Prof. Mohamed Neffati, an expert in the field of the ethnobotanical value of the spontaneous plants of the Tunisian arid zones, for botanical identification.

